# Polyanionic peptibody ameliorates tau pathology and promotes neuronal proteostasis in a tau transgenic mouse model

**DOI:** 10.1101/2025.08.13.665777

**Authors:** Alayna C. Caruso, Tomas Kavanagh, Ashley J. van Waardenberg, Mark E. Graham, Eleanor Drummond, Rebecca M. Nisbet

## Abstract

The intraneuronal aggregation of tau is a key driver of pathogenesis in Alzheimer’s disease and other tauopathies. The positively charged proline-rich region and microtubule-binding domain of tau are important for tubulin binding which enables axonal transport in physiological conditions. In disease, however, the positively charged repeat domains of such regions become exposed, facilitating tau-tau aggregation and other pathogenic tau-protein interactions which leads to neuronal dysfunction. Here, we generated a polyanionic peptide to electrostatically interact with tau and fused it to an inert single-chain variable fragment (scFv) antibody for improved peptide half-life and downstream intraneuronal expression. The resultant polyanionic peptibody, ACR12, was investigated *in vitro* and in tau transgenic mice for ability to prevent tau aggregation and associated disease phenotypes. In human neuroblastoma cells, ACR12 was stably expressed within the cytoplasm where it successfully engaged intracellular tau. DNA encoding ACR12 was then packaged into a brain-penetrant Adeno-associated virus, AAV-PHP.eB, for intravenous delivery into K3 tau transgenic mice. Such treatment facilitated long-term intraneuronal expression of ACR12 and reduced total and phosphorylated tau in the brain. ACR12 treatment in female K3 mice also restored deregulated neuronal proteins to wild-type levels. This study highlights the role of positively charged domains in tau pathogenesis and demonstrates the therapeutic utility of electrostatic tau interactors.

## INTRODUCTION

Tauopathies are progressive neurodegenerative diseases characterized by the abnormal aggregation and deposition of the tau protein. Tau comprises of multiple functional domains including two positively charged regions: the proline-rich region (PRR) and the microtubule-binding domain (MTBD). In its native state, the positively charged residues throughout these regions enable tubulin binding for regular axonal transport, and contribute to intermolecular repulsion between tau molecules, helping to maintain tau in a soluble, non-aggregated conformation^1–5^. In tauopathies, however, the dysregulation and modification (e.g., hyperphosphorylation) of tau alter its charge distribution and overall conformation, weakening the association of tau with microtubules, disrupting tau-protein interactions, mislocalising tau, and promoting the pathological self-assembly of tau into aggregates or neurofibrillary tangles (NFTs)^4,6–14^. The aggregation of tau is primarily driven by the PRR and MTBD^15–17^, and occurs in stages, beginning from native state monomeric tau that misfolds into dimers, followed by soluble oligomers, protofibrils, and finally insoluble fibrils and NFTs^2^. Whilst NFT accumulation in the brain is associated with disease progression^18,19^, tau oligomers have been demonstrated to be most neurotoxic, contributing to synaptic dysfunction^20–22^.

Beyond having roles in aggregation, the PRR and MTBD also mediate interactions of tau with other macromolecules and cellular structures, including proteins such as kinases, membrane receptors, RNA, and lipids^7,8,22–34^. These interactions are strongly influenced by tau phosphorylation and thus, in disease become dysregulated to cause downstream neuronal dysfunction^35^. Interestingly, most phosphorylation sites found across primary tauopathies cluster within the PRR, underscoring the relevance of this region in disease^8^. One example of such dysregulation is the electrostatic interaction of tau with RNA and RNA-binding proteins^8,22,25,26^. Under cellular stress, RNA-binding proteins mislocalise to the cytoplasm and bind tau oligomers, promoting stress granule formation that impairs protein translation and synaptic functioning^22,35,36^. Another example is tau’s electrostatic interaction with heparin sulfate proteoglycans (HSPGs) which function as cell surface receptors to mediate the internalisation of tau ‘seeds’, contributing to the transcellular spread of tau pathology^27–29^. Similarly, negatively charged lipid headgroups of cell membranes directly interact with the PRR and MTBD of tau, promoting tau aggregation and propagation^30–34^. Evidently, the positively charged amino acid stretches of tau and associated electrostatic interactions contribute to cellular dysfunction in disease, making the mid-region of tau a promising target for therapeutic intervention.

Here, we endeavoured to inhibit tau pathogenesis using an engineered polyanionic peptide that targets the positively charged PRR and MTBD of tau. As peptides have a short half-life and are ineffective at crossing the neuronal plasma membrane, we fused the peptide to an inert single-chain variable fragment (scFv) antibody to facilitate intraneuronal expression and aid in sterically hindering pathogenic tau interactions. The resulting peptibody, ACR12, was demonstrated to bind tau *in vitro,* as well as intracellularly following expression in human neuroblastoma cells. Intravenous administration of the ACR12 gene packaged into a brain-penetrant Adeno-associated virus (AAV) serotype, AAV-PHP.eB, led to a significant reduction of total and phosphorylated tau levels in K3 mice relative to Sham controls. Furthermore, ACR12 treatment restored the K3 mouse brain proteome towards a wild-type-like profile, suggesting rescue of proteostasis.

## RESULTS

### ACR12 peptibody binds recombinant tau *in vitro*

Three polyanionic peptides were generated with increasing net negative charge of −3.4, −9.4, and −12.6, and fused to a non-specific scFv to form peptibodies, ACR3, ACR9, and ACR12 (Figure 1A). The PRR and MTBD of tau comprise of positively charged repeats, making these regions favourable to electrostatic interaction with polyanions (Figure 1B). To investigate this, an ELISA was performed to assess binding of recombinant ACR proteins to human full-length tau (hTau-441), in which ACR12 demonstrated the greatest binding capacity, followed by ACR9. ACR3, however, did not bind tau at all, suggesting that ACR/tau binding is charge dependent (Figure 1C). Drug potency was determined via EC_50_ calculation to be 12.41uM for ACR9 and 2.46uM for ACR12 (comparison of ACR9 and 12 via *F-*test, P<0.0001) (Figure 1D). ACR3 showed no detectable interaction with recombinant hTau-441, potency was therefore unable to be determined, and this construct was not explored further. A subsequent ELISA was performed to test interaction with tau’s MTBD using the K18 tau fragment. This test revealed comparable binding (P=0.400) and potency (P=0.436) of ACR9 and ACR12 with the MTBD of tau (Figure 1E and F).

**Figure 1.**
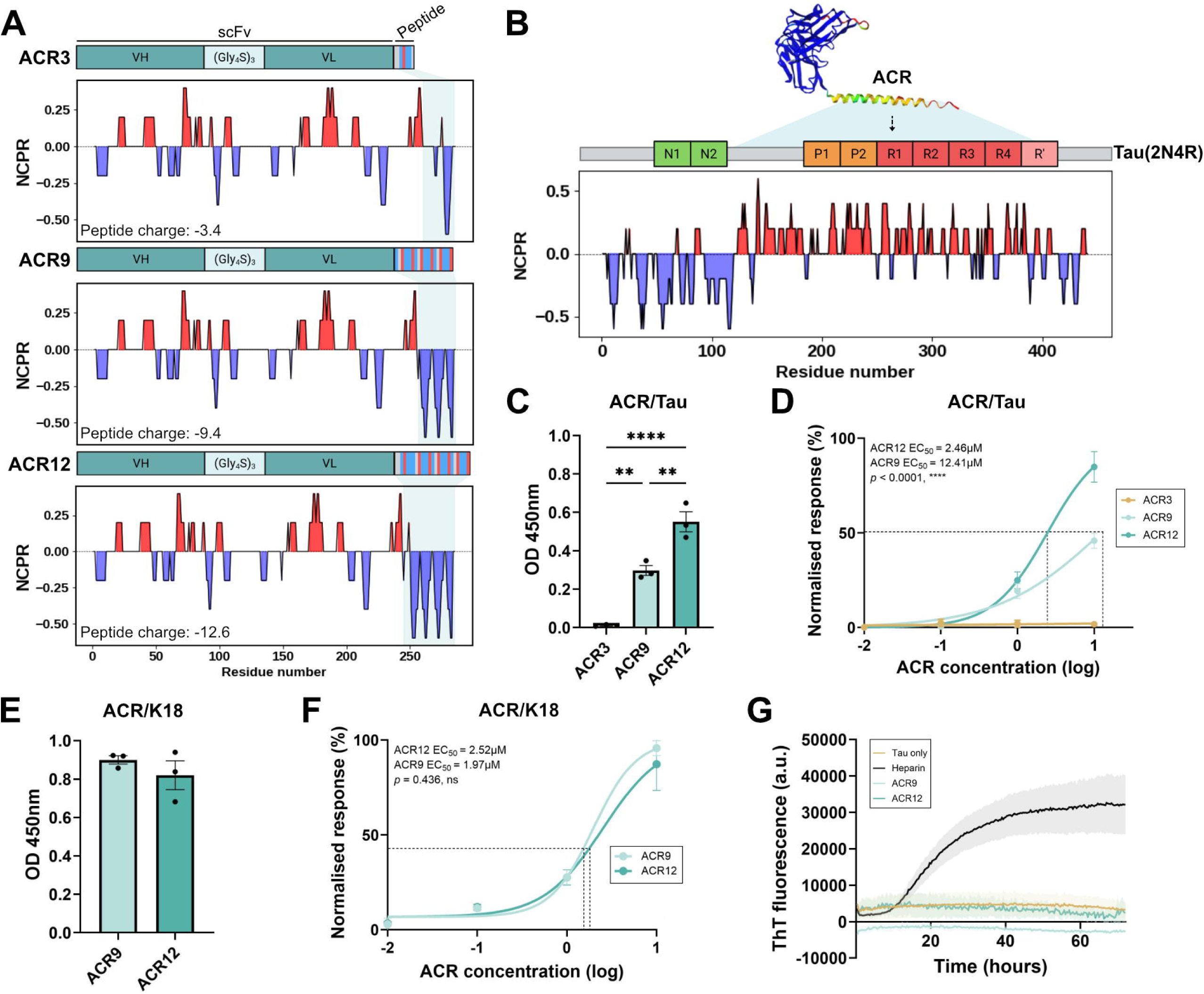
ACR9 and ACR12 bind recombinant tau in a charge-dependent manner. (**A**) Schematic representation of peptibodies, ACR3, ACR9, and ACR12, with net charge per residue (NPCR) distribution. (**B**) Schematic of predicted ACR interaction with positively charged regions of full-length human tau (hTau-441; 2N4R), AlphaFold-predicted ACR12 structure aligned with tau NPCR. (**C**) ELISA of ACR3, ACR9, and ACR12 at maximum dose (10uM) with recombinant hTau-441, analysed via one-way ANOVA with Tukey’s multiple comparisons (** = *p* < 0.01; **** = *p* < 0.0001). (**D**) Determination of ACR concentration yielding half-maximal binding (EC_50_) to hTau-441 by ELISA; No EC_50_ was determinable for ACR3 due to low binding; ACR9 and ACR12 EC_50_ compared via *F*-test (****, *p* < 0.0001). (**E**) ELISA of ACR9 and ACR12 at maximum dose with recombinant tau microtubule-binding domain (K18 fragment; tau-K18), analysed via Welch’s unpaired *t*-test (ns, *p* = 0.3999). (**F**) EC_50_ of ACR9 and ACR12 with tau-K18 by ELISA; curved compared via *F*-test (ns, *p* = 0.436). (**G**) Recombinant tau fibrillization measured overtime via ThT fluorescence in samples of hTau-441 incubated alone or with ACR9, ACR12, or heparin.

Linear polyanions, such as heparin and RNA, are well-established to induce tau fibrillization^37–39^. To ensure ACR peptibodies do not scaffold tau aggregation in the same way, thioflavin T (ThT) fluorescence of recombinant hTau441/ACR preparations was measured. Unlike heparin, ACR12 and ACR9 did not induce tau fibrillization (Figure 1G).

### ACR12 engages with cytoplasmic tau in human neuroblastoma cells

Transfection of SH-SY5Y human neuroblastoma cells that overexpress GFP-tagged full-length human tau (Tau-GFP cells) with ACR9 or ACR12 yielded comparable peptibody expression (Figure 2A and B). Immunofluorescent microscopy revealed that within the cell, both ACR9 and ACR12 localised to the cytoplasm and nucleus, in close proximity to cytoplasmic tau (Figure 1C and D). More specifically, an average Pearson’s coefficient of 0.37 for ACR9 and 0.45 for ACR12, was calculated, suggesting spatial overlap of ACR9 and ACR12 with tau (Figure 1D). To investigate a direct interaction between ACR9 and ACR12 with tau, a GFP pull-down assay was performed on ACR9- and ACR12-transfected Tau-GFP cells. ACR12 and, to a lesser extent ACR9, was detected in the pull-down sample, confirming engagement with intracellular tau (Figure 2E).

**Figure 2.**
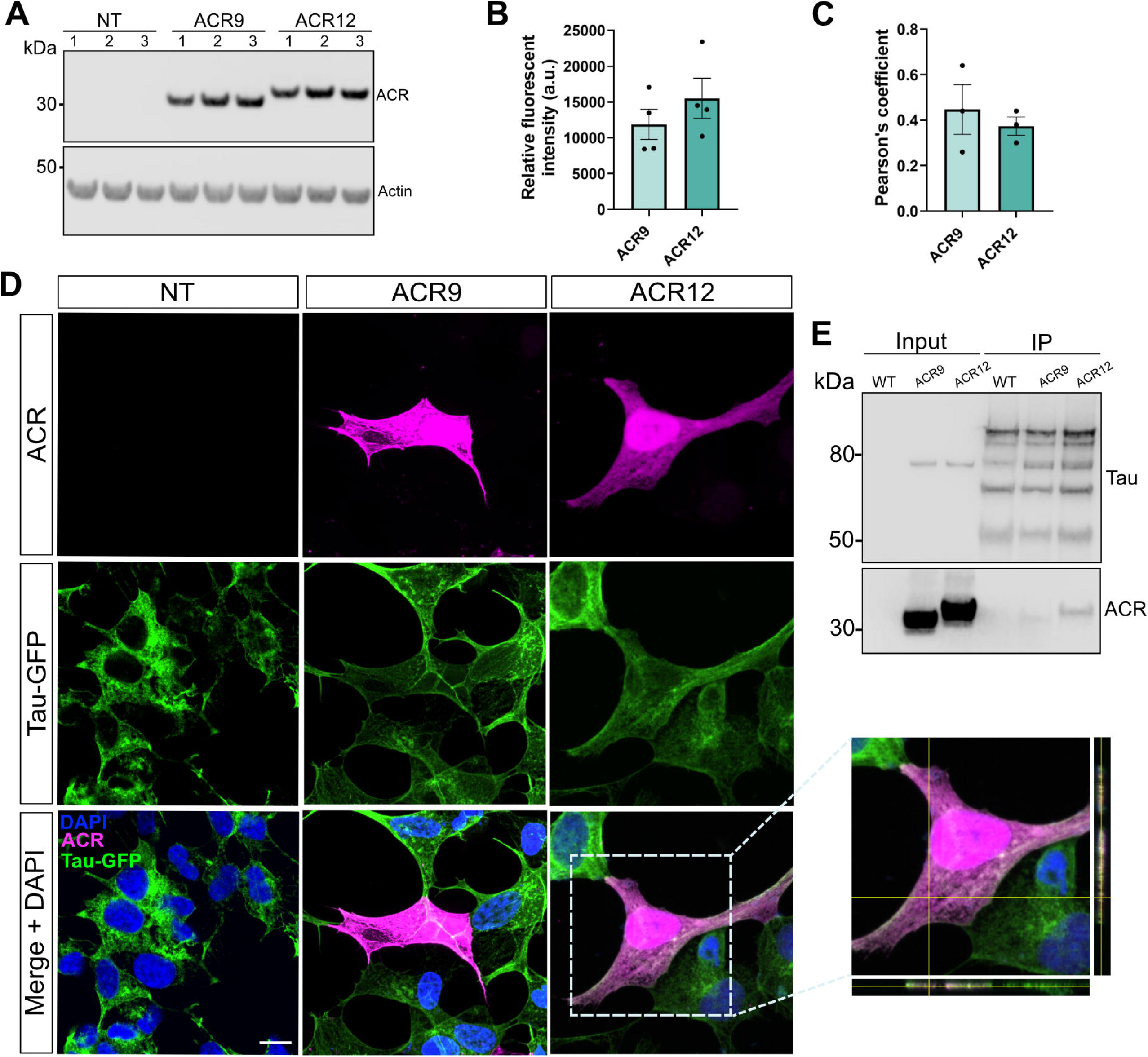
ACR12 is expressed within human neuroblastoma cells and engages intracellular tau. (**A**) Immunoblot of non-transfected (NT) Tau-GFP cells or cells transfected with ACR9 or ACR12, probed with α-Flag for ACR detection and β-actin. (**B**) Quantification of *B*, points normalised to β-actin. (**C**) Pearson’s correlation coefficient between tau and ACR proteins expressed in Tau-GFP cells. (**D**) Representative images (60x) of NT, ACR9, or ACR12 Tau-GFP cells. Cell nuclei stained with DAPI (blue), ACR labelled with α-Flag (magenta), tau detected with GFP (green); Scale bar = 10μm. (**E**) Immunoblot of non-tau-expressing WT SH-SY5Y cells, ACR9-, or ACR12-transfected Tau-GFP cells before (input) and after GFP pull down (IP), probed with α-Flag or Tau-5. Three-four independent experiments were performed for each assay. Data (B and C) analysed by Welch’s unpaired t-test (ns, *p* = 0.342 and 0.583, respectively).

### Intravenous AAV-mediated delivery of ACR12 facilitates neuronal transduction in wild-type and K3 tau transgenic mice

To investigate intraneuronal expression of ACR12 *in vivo*, the DNA encoding ACR12 was packaged into the brain-penetrant AAV-PHP.eB and injected via the tail vein into five-week-old K3 tau transgenic mice. ACR9 was also packaged into AAV-PHP.eB and included as an intermediate-charge comparator. The AAV-PHP.eB serotype was selected as it has been shown to facilitate efficient brain transduction in C57BL/6J mice following intravenous (IV) injection^40^. K3 mice overexpress human tau with a K369I mutation and demonstrate a progressive accumulation of tau inclusions^41,42^. We also explored IV injection of AAV-PHP.eB ACR12 in wild-type (WT) mice. Two-weeks post-treatment, both ACR9 and ACR12 could be detected in the K3 brain at comparable levels, and no expression was observed in the peripheral organs (Figure 2A). ACR12 was also detected in the WT brain at a comparable level to what was observed in K3 mice, demonstrating that overexpression of human mutant tau does not affect AAV neuronal transduction (Figure 2A). ACR12 expression in the K3 and WT brain was also detected three-months post-injection, suggesting that a single IV administration may be sufficient for long-term treatment (Figure 2B). Immunofluorescent microscopy revealed neuronal ACR9 and ACR12 expression across multiple brain regions, including the cortex, thalamus, and hippocampus (Figure 2C and D).

### Treatment with ACRs is well-tolerated but does not improve motor function in K3 mice

To investigate the therapeutic effect of ACR12 treatment, a new cohort of K3 mice at five-weeks of age were intravenously injected with saline (Sham), AAV-PHP.eB-packaged ACR9, or AAV-PHP.eB ACR12 (Figure 3A). K3 mice characteristically display lower weight, as well as a motor impairment from birth when compared to WT mice^41,42^. Weight was recorded weekly and behavioural tests were conducted to assess motor function. Non-transgenic WT littermates were used as healthy controls for all behavioural analyses. No difference in weight between ACR9 or ACR12 treated compared to Sham K3 mice was observed (Figure 3B). Fortnightly rotarod testing showed no difference in latency to fall recorded for ACR9 or ACR12 treated mice when compared to Sham mice (Figure 3C). At endpoint, no differences were found in digigait recorded stride length between ACR9 and ACR12 treated mice compared to Sham controls (Figure 3D). Taken together, whilst ACR treatment did not improve weight gain nor motor deficits in K3 mice, treatment did not cause observable adverse effects.

**Figure 3.**
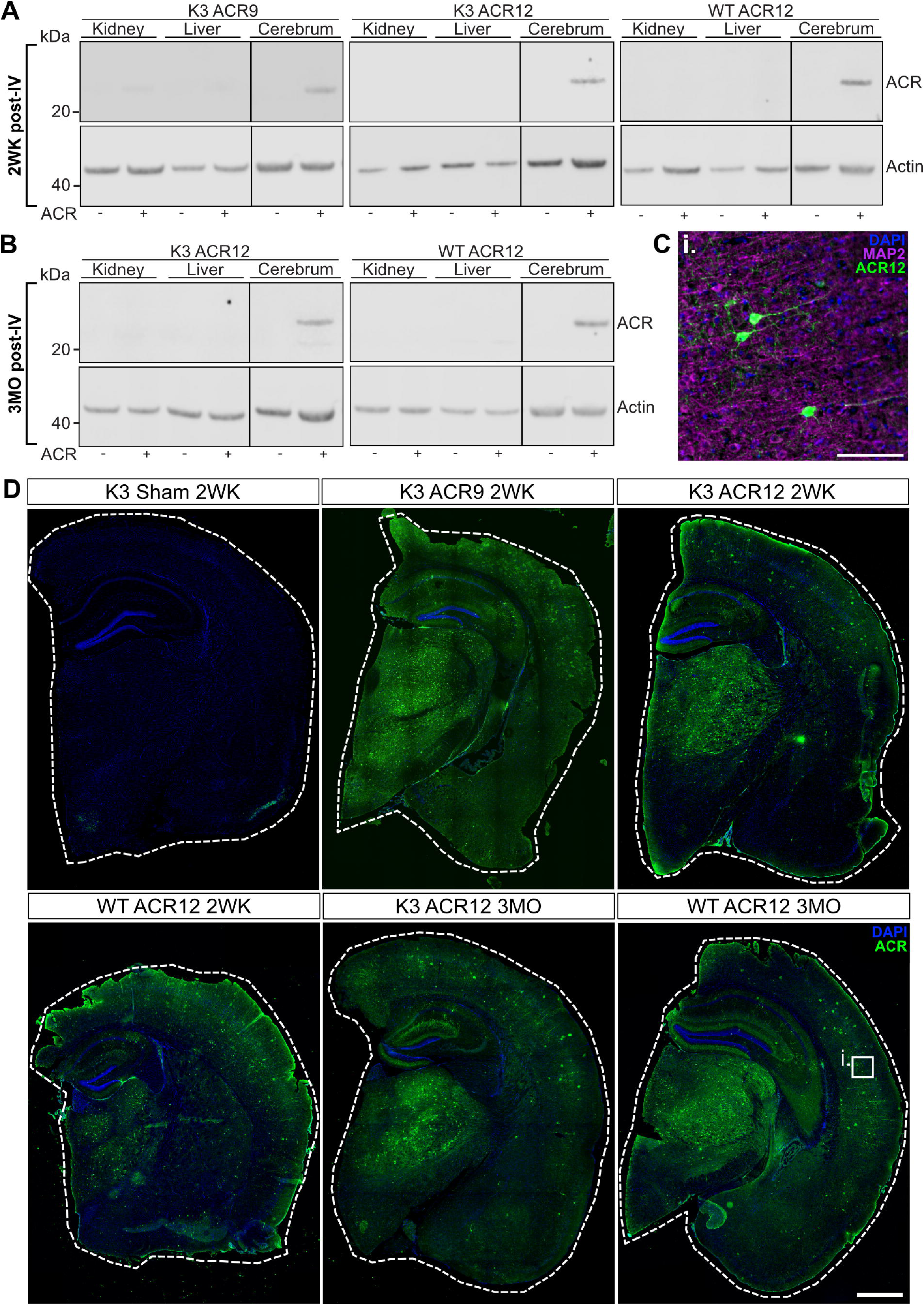
Intravenous AAV-mediated delivery of ACRs in K3 and wild-type mice facilitates intraneuronal transduction. (**A**) Immunoblots of tissue homogenates collected from K3 or wild-type (WT) mice two-weeks following intravenous (IV) injection with saline (Sham; scFv -) or AAV-PHP.eB-packaged ACR9 or 12 (scFv +). (**B**) Immunoblots of tissue homogenates collected from ACR12-treated mice three-months post-injection. All immunoblots probed with α-Flag and β-actin. (**C**) Representative image of cortical neurons expressing ACR12 (scale bar = 100μm). (**D**) Representative images (20x) of coronal half-brain sections collected from Sham and ACR-treated mice, two-weeks or three-months post-injection (scale bar = 1000μm). For immunofluorescent microscopy, cell nuclei stained with DAPI (blue), ACR with α-Flag (green), and neurons with MAP2 (magenta).

### ACR12 treatment significantly reduces total and phosphorylated tau in K3 tau mice

Following behavioural analysis, the effect of ACR12 treatment on pathological tau accumulation and phosphorylation was assessed. Total tau levels in the brain were significantly reduced by 25.71% in K3 mice treated with ACR12 when compared to Sham controls and ACR9-treated mice (P < 0.0001) (Figure 4A and B). This effect was more severe in female K3 mice (39.38% reduction compared to Sham; P < 0.0001), than in males where only a trend in reduction was observed (9.12%; P = 0.2164) (Figure 4B). ACR9 had no effect on total tau levels in K3 mice compared to Sham (Figure 4A and B).

**Figure 4.**
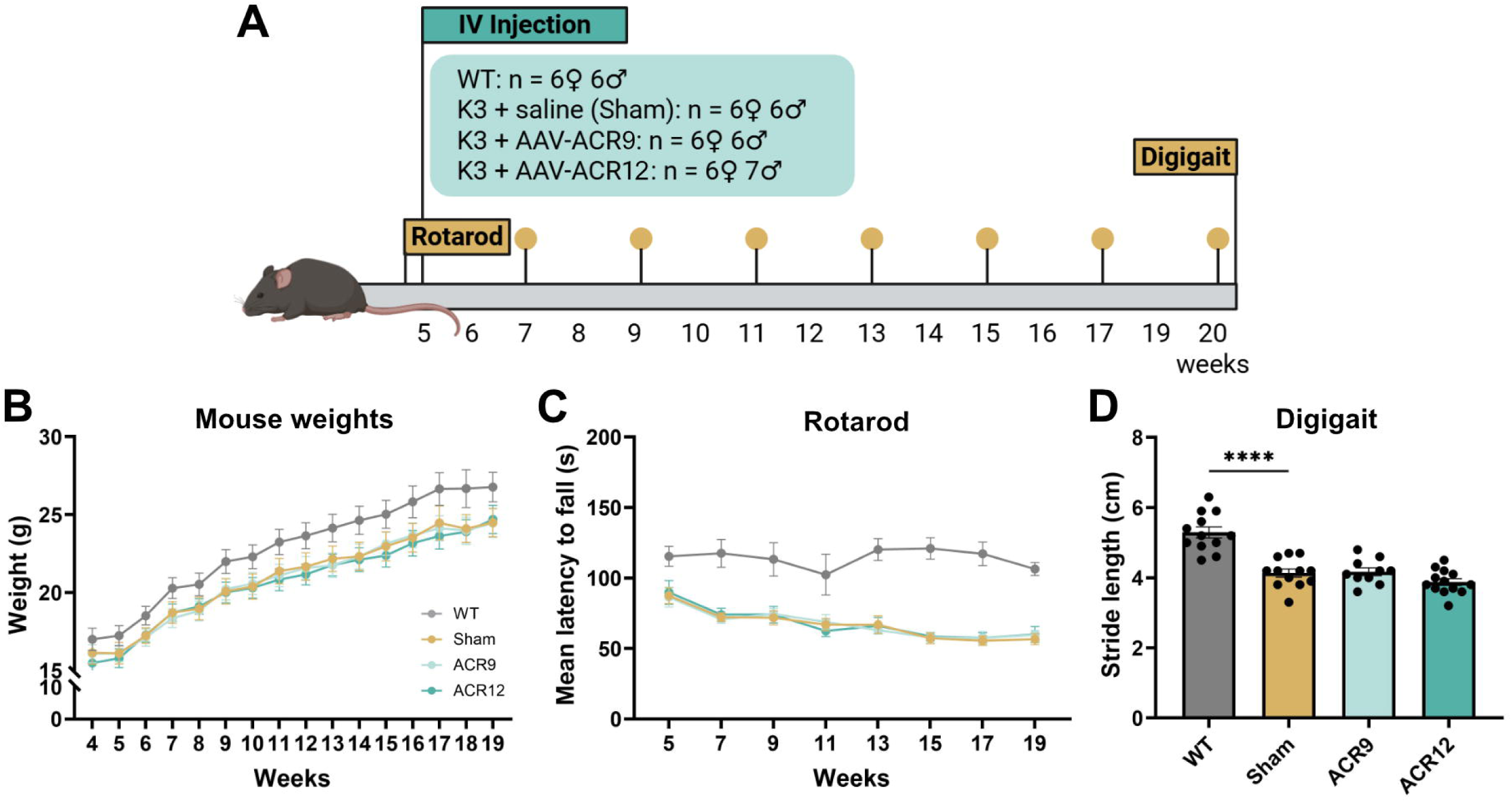
ACR9 and ACR12 treatment is well-tolerated but does not improve motor deficits in K3 tau transgenic mice. (**A**) Treatment schedule with behaviour: at five-weeks-old mice were injected intravenously with saline (Sham), AAV-PHP.eB ACR9, or ACR12. Following injection, mice underwent biweekly Rotarod and endpoint Digigait testing. (**B**) Mean weight of treatment groups over time. (**C**) Mean latency to fall from Rotarod for treatment groups over time. (**D**) Stride length recorded via Digigait. All data (B, C, and D) analysed using one-way ANOVA with Tukey’s multiple comparisons (**** = *p*<0.0001).

Phosphorylated tau (p-tau) pathology was quantified via immunofluorescent microscopy using antibodies specific for p-tau epitopes: AT180 (Thr231), AT8 (Ser202/Thr205), and RN235 (Ser235) (Figure 4C-H). Importantly, tau phosphorylated at Thr231 (AT180) is an early marker of tauopathy, whilst tau phosphorylated at Ser202/Thr205 (AT8) and Ser235 (RN235) are associated with late-stage disease^43,44^. ACR9-treated K3 mice showed a trend in reducing cortical AT180 p-tau in comparison to Sham (31.34%; *P* = 0.1203), which was significant for female ACR9-treated mice only (76.5%; *P* = 0.0213). ACR12 on the other hand, facilitated a significant 62.5% reduction in AT180 p-tau compared to Sham (*P* = 0.0012). This effect that was more robust in female mice (45.62%; *P* = 0.0016) than males (40.19%; *P* = 0.0609) (Figure 4F). When comparing Sham female and male K3 mice, females had a higher AT180 load (39.58% more; *P* = 0.0056), possibly accounting for the enhanced ACR-mediated effect observed in treated female mice compared to males. For AT8 p-tau, ACR9 treatment only showed a minor trend in reduction compared to Sham (17.48%; *P* = 0.6798). Whereas ACR12 reduced AT8 p-tau by 52.19% compared to Sham controls (*P* = 0.0324). When separating the data by sex, however, only a trend in AT8 reduction was achieved by ACR12 for males (40.61%; *P* = 0.3853) and females (66.01%; *P* = 0.1027) (Figure 4G). For RN235, ACR9 again only exhibited a slight trending reduction compared to Sham (36.08%; *P* = 0.1523). Whilst ACR12 treatment induced a significant 54.28% reduction in RN235 p-tau (*P* = 0.0189). As RN235 detects tau neurofibrillary tangles^43^, this result suggests a role for ACR12 in preventing pathological tau aggregation. Interestingly, ACR12 male mice showed a more significant reduction (64.41%; *P* = 0.0418) than treated female mice (33.67%; *P* = 0.3385) (Figure 4H). Although of note, Sham K3 males showed a trended increase in RN235 p-tau when compared to Sham K3 females (50.87%; P = 0.0773). Together, these results suggest that ACR12 is effective in reducing total and phosphorylated tau load in the brains of K3 mice. This therapeutic effect was observably greater than that of ACR9, likely due to its more negative charge. As such, we continued on with ACR12 alone for further study.

### ACR12 treatment promotes neuronal proteostasis in female K3 tau mice toward wild-type levels

Tau plays a critical role in neuronal processes and, as such, widespread changes in proteostasis have been reported in human tauopathy brain tissue and in cell and animal models of disease ^7,8,45,46^. Quantitative proteomics was therefore conducted to assess the impact of ACR12 treatment on the K3 mouse proteome and to investigate its ability to restore proteomic dysregulation specifically related to neuronal function considering it is expressed within neurons. First, the K3 and WT mouse proteomes were compared to identify the proteomic alterations that resulted from human tau K369I overexpression and pathogenic tau accumulation in our mouse model. Such analysis confirmed that K3 mice have a significantly different proteome from WT mice, with a more pronounced effect observed in females (Figure 5A and B). Specifically, K3 mice exhibited 1,656 and 1,007 differentially expressed proteins (DEPs; abundance fold-change *P* < 0.05 after FDR correction) when compared to WT female and male mice, respectively. These DEPs were considered to be associated with K369I tau pathology. Of these proteins, 917 were decreased and 739 increased in female K3s (Figure 5A), and 585 were decreased and 422 increased in males (Figure 5B). Interestingly, only 35.2% of DEPs increased in female K3 mice compared to WT were also increased in males (Figure 5C). Similarly, only 29% of DEPs that decreased in female K3 mice were shared with male K3 mice (Figure 5C). This suggests that each sex has a unique proteomic signature, when comparing K3 to WT mice.

**Figure 5.**
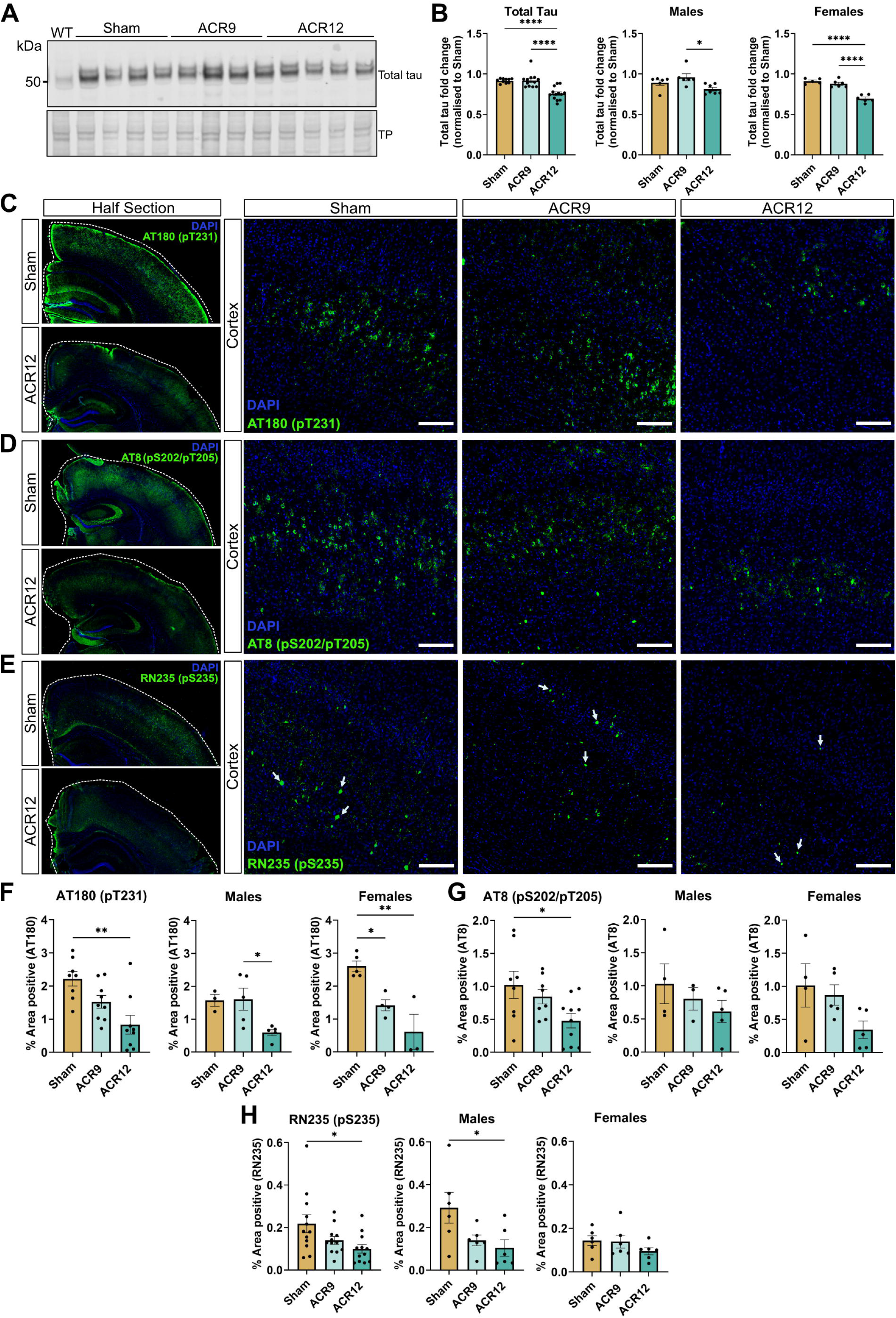
ACR12 reduces total and phosphorylated tau in K3 tau transgenic mice. (**A**) Immunoblot of total brain homogenates collected from WT, K3 Sham, K3 ACR9, and K3 ACR12 mice, probed with Tau-5; total protein (TP) detected with REVERT. (**B**) Quantification of *(A)*, data reported as fold change to Sham; representative of three-four independent experiments (n = 12-13 per treatment group). (**C**) Representative images (20x) of half-brain sections and cortical tissue (scale bar = 100μm) collected from Sham and treated K3 mice. Cell nuclei stained with DAPI (blue), and tau phosphorylated at Thr231 labelled with AT180 (green) or (**D**) tau phosphorylated at Ser202/Thr205 labelled with AT8 (green) or (**E**) tau phosphorylated at Ser235 with RN235 (green). (**F**) Quantification of AT180-positive cortical area in Sham and treated K3 mice, presented as percent area positive (n = 8-9 per treatment group). (**G**) Quantification of AT8-positive cortical area (n = 8-10 per group). (**H**) Quantification of RN235-positive cortical area (n = 12 per treatment group). Immunofluorescent microscopy is representative of two-three sections per mouse. All data (B, F, G, and H) analysed via one-way ANOVA with Tukey’s multiple comparisons (* = p<0.05; ** = p<0.01; **** = p<0.0001).

Principal component analysis (PCA) revealed that ACR12-treated female K3 mice formed a distinct cluster that shifted away from K3 Sham and towards WT (Figure 5D). This was also observed in males (Figure 5E), suggesting a restorative effect of ACR12 on proteostasis (Figure 5F). Quantification of proteins in ACR12-treated K3 mice compared to K3 Sham mice revealed that treatment significantly altered the female K3 proteome. Specifically, 538 DEPs, of which 300 decreased and 238 increased, were observed in the ACR12-treated female K3 brain (Figure 5G). A much smaller response was observed in the male ACR12-treated K3 mice, wherein only 31 DEPs were observed, of which 11 decreased and 20 increased in abundance (Figure 5H). This indicates that ACR12 had a more pronounced effect in female K3 mice than in males and, given the observed sex differences in the K3 mouse proteome, subsequent proteomic analyses focused exclusively on female K3 mice.

To assess the ability of ACR12 to reverse disease-associated proteomic alterations, the abundance of the 1,656 DEPs identified in K3 mice when compared to WT mice (see Figure 5A) were investigated in female ACR12-treated K3 mice. ACR12 increased the abundance of the majority of proteins that were downregulated in K3 relative to WT (789 of 917; Quadrant I) and decreased the levels of most proteins that were upregulated in K3 relative to WT (623 of 739; Quadrant III) (Figure 5I). In total, 1,412 of 1,656 DEPs (85.2%) shifted towards WT following treatment with ACR2, suggesting ACR12 may restore WT-like proteostasis in female K3 mice (Figure 5I). Of the 1,412 proteins, 242 were identified to be significantly altered in both K3vWT and K3 ACR12vK3 Sham comparisons, representing disease-associated proteins that were significantly impacted by treatment (Figure 5I). Linear regression modelling confirmed a strong ACR12-mediated reversion towards WT levels among these 242 proteins (*R*^2^ *=* 0.92, *P* = 5.7 × 10 ¹³²; Figure 5I). Comparison of the abundance of these proteins across K3 Sham, K3 ACR12, and WT groups, revealed a similar expression profile between K3 ACR12 and WT groups (Figure 5J). Specifically, 238 of the 242 proteins (98.4%) reverted toward WT levels following treatment (Figure 5J). Confirming ACR12s reversion effect, a positive correlation between changes in protein levels in ACR12-treated K3 mice and WT mice relative to K3 Sham was determined (Pearson’s correlation *r* = 0.96, *P =* 5.74 × 10 ¹³²). This result validates that ACR12 treatment induces proteomic reversion of pathology-affected proteins toward WT levels.

To determine which cell types the reverted proteins were predominantly expressed within, we leveraged brain single-cell data from the Human Protein Atlas^47^ to determine cellular specificity across 19 primary cell types. The collective gene set encoding the reverted proteins demonstrated neuronal specificity, with dominant expression in excitatory, inhibitory and other neurons, followed by oligodendrocytes and oligodendrocyte progenitor cells (OPCs) (Figure 5K). While genes encoding these proteins were consistently expressed at the highest levels in neurons, expression was also observed in astrocytes, vascular endothelial cells, and microglia (Figure 5L). These findings suggest that ACR12s reversion effect predominantly impacts neuronal cell types.

### ACR12 treatment restores proteins important for neuronal functions

Pathway enrichment analysis of the 238 disease-associated proteins that reverted with ACR12 treatment (identified in Figure 5J) showed that ACR12-upregulated proteins were largely enriched in pathways including RNA splicing, mRNA processing, metabolic process, and cellular organisation (Figure 6A). Protein-protein interaction (PPI) and subsequent Molecular Complex Detection (MCODE) analysis of proteins associated with the top 10 increased pathways (by FDR) identified two densely connected clusters of proteins linked to RNA processing (FDR = 5.98X10^-9^) and synaptogenesis and neuronal membrane trafficking (FDR = 0.0118) (Figure 6B). RNA splicing is dysfunctional in tauopathy ^8,48^, and several RNA-binding proteins with important roles in RNA splicing and related mRNA/RNA processing were identified to be increased towards WT levels in ACR12-treated K3s, for example, hnRNP A1, hnRNP C, hnRNP F, RBM11, TAF15, and YBX1 (Figure 6B). Additionally, proteins with roles in regulation of neuronal plasticity (CRTC1 and DCC) and synaptic processes (ANK3, SEPT8, SEPT11, STXBP1, NRXN3) were also restored in female K3 mice treated with ACR12 (Figure 6B). Together, these findings indicate that ACR12 treatment may restore RNA splicing and related RNA processes, whilst regulating neuronal and synaptic organisation and plasticity.

**Figure 6.**
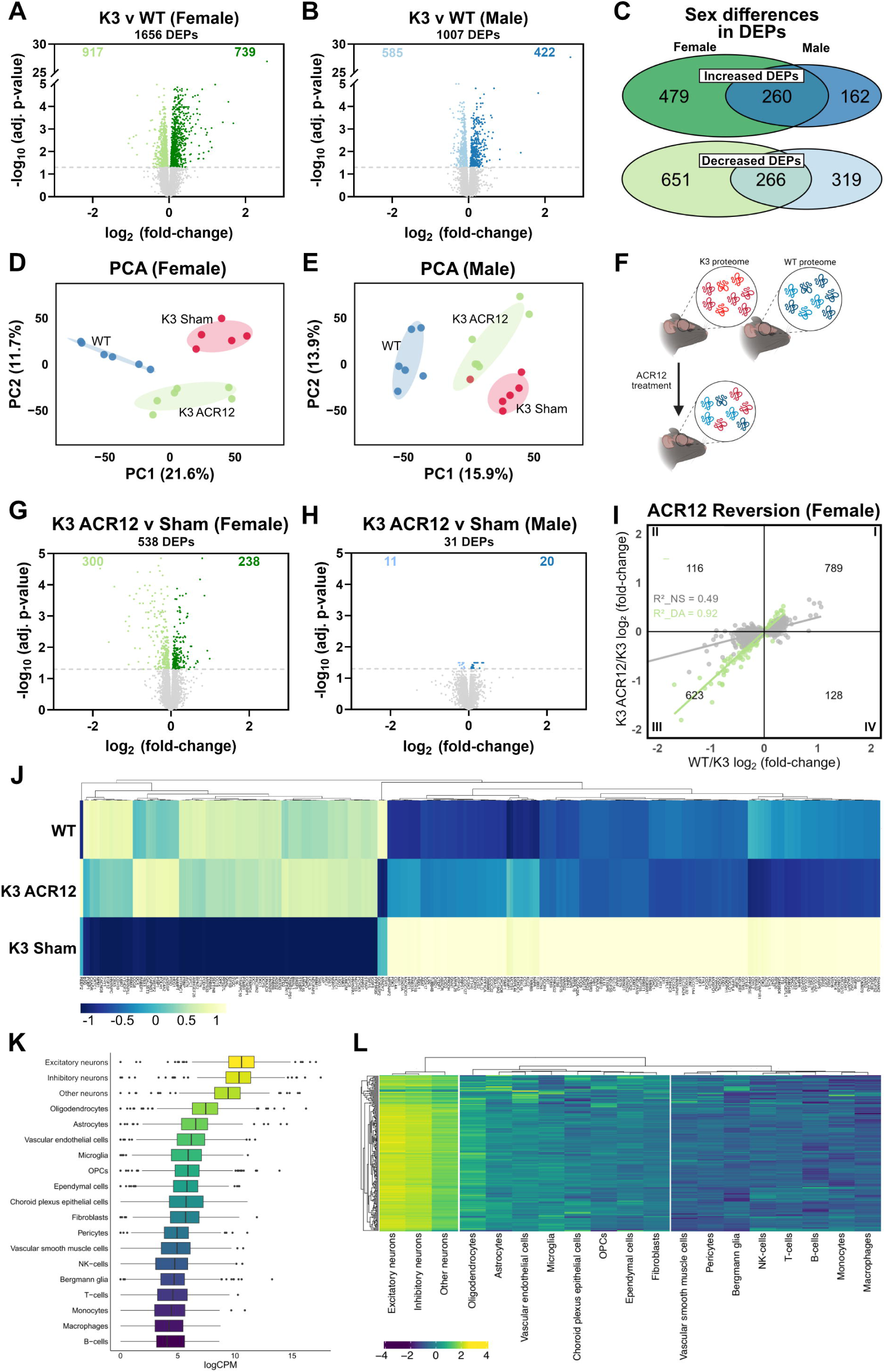
ACR12 restores proteomic deregulation in female K3 tau transgenic mice. (**A**) Volcano plot showing DEPs in K3 female, and (**B**) K3 male mice relative to WT. (**C**) Venn diagram of intersecting DEPs in female and male K3 mice relative to WT. (**D**) PCA of female mice for PC1 and PC2 revealing three clusters: WT mice, K3 Sham, and ACR12-treated K3s, wherein K3 ACR12 shifts towards WT. (**E**) PCA of male mice. (**F**) Schematic representation of the ACR12-mediated shift of the K3 brain proteome towards WT. (**G**) Volcano plot of DEPs in female, and (**H**) male K3 ACR12 mice relative to Sham. (**I**) Reversion plot displaying the shift of DEPs identified in *(A)* in ACR12-treated female K3s. Quadrant I and III hold proteins that shifted towards WT with treatment, QI proteins increased and QIII decreased in WT and ACR12 K3 compared to Sham K3. Grey proteins (1,412 of 1,656) were significantly altered in K3vWT but only trended change in K3 ACR12vK3 Sham; Green proteins were significantly altered in both comparisons (242 of 1,656), representing disease-associated proteins that significantly changed with treatment. Reversion analysed via simple linear regression (*R*^2^ = 0.92) and significance of slope assessed by two-tailed unpaired *t*-test (*P* = 5.7 × 10 ¹³²). (**J**) Heatmap illustrating z-scores of log_2_ transformed abundances of 242 proteins identified in *(I),* quantified across groups. 238 of 242 proteins (98%) reverted to WT levels with ACR12-treatment; correlation between K3 ACR12 and WT protein abundances analysed via Pearson’s correlation with two-tailed *t-*test (*r* = 0.96, *P =* 5.74 × 10 ¹³²). (**K**) Boxplots representing the distribution of cell-specific expression of genes encoding ACR12-reverted proteins identified in panel *(J).* LogCPM represents log2 transformed normalised counts per million per gene identified in the single-cell RNA-seq data. Cell type (y-axis) ordered highest (top) to lowest (bottom) median expression of the gene set. (**L**) Heatmap illustrating z-scores of log2 transformed abundances of single-cell expression profiles of reverted protein genes. Cell types grouped in 3 clusters (tree dendrogram cut at k=3, grey dotted line).

**Figure 7.**
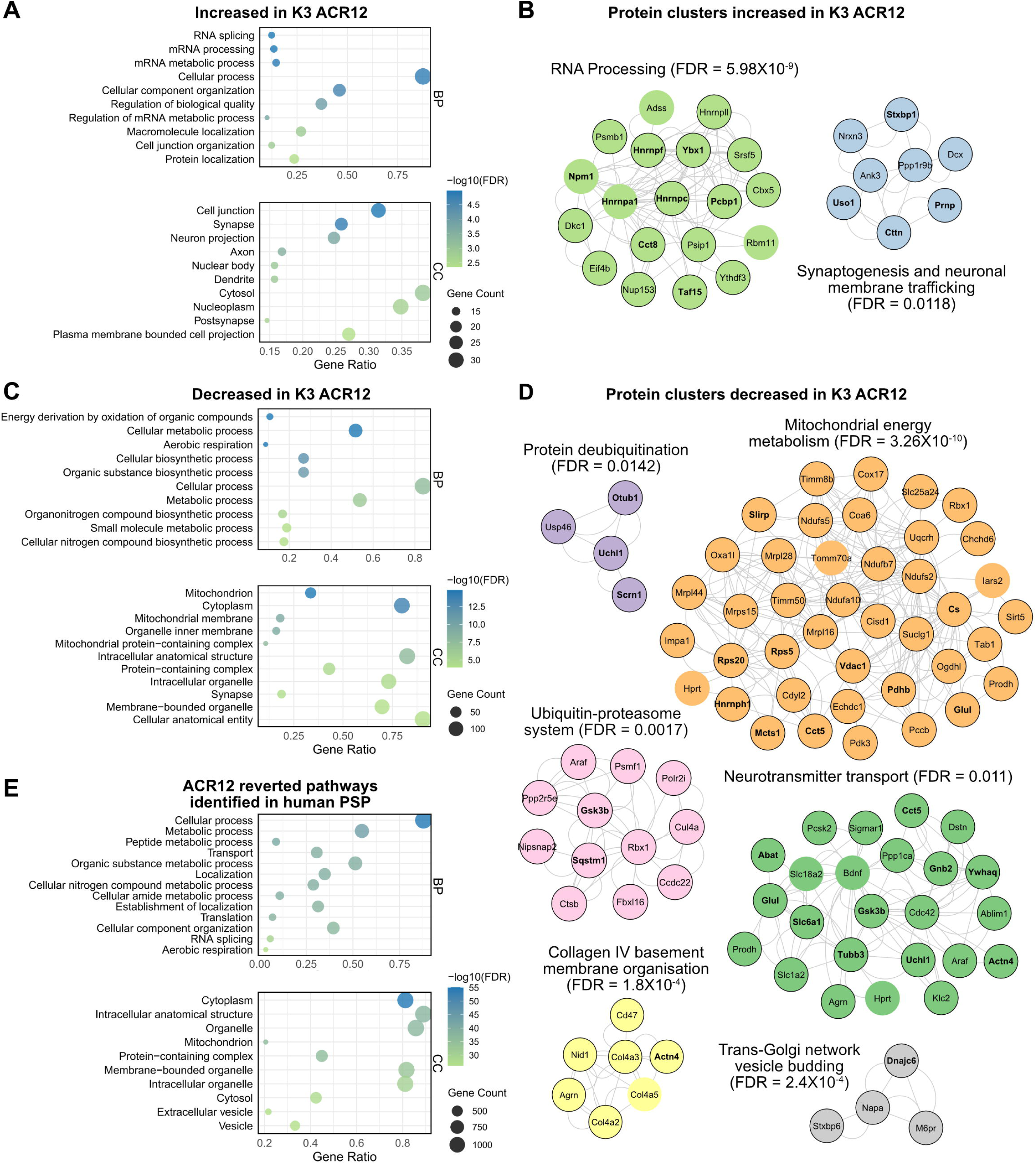
ACR12 restores proteins important for neuronal functions in female K3 mice. (**A**) Top 10 (ordered by FDR) enriched biological processes (BP) and cellular components (CC) found to be increased in K3 ACR12 female mice towards WT levels. (**B**) Protein interaction clusters identified via MCODE analysis of top 10 pathway proteins that increased with ACR12 treatment in K3 females. Bolded proteins are known tau interactors^8^, and protein nodes with black border represent proteins matched to human orthologs in a PSP protein dataset^51^. (**C**) Top 10 BP and CC terms that decreased in ACR12-treated K3 female mice towards WT. (**D**) MCODE clusters of top 10 decreased pathway-associated proteins. (**E**) Top 10 BP and CC terms identified from the 1,319 orthologous proteins that were significantly altered with ACR12 treatment in female K3 mice and found in a human PSP protein dataset^51^. Cellular component organisation, RNA splicing, and aerobic respiration additionally included due to biological relevance (also significant by FDR).

Proteins that downregulated towards WT levels with ACR12 treatment were enriched in pathways linked to mitochondrial and cellular respiration functions, for example, energy derivation by oxidation of organic compounds, cellular metabolic process, and aerobic respiration (Figure 6C). PPI and MCODE analyses of the top 10 decreased pathway-associated proteins revealed six protein clusters linked to mitochondrial energy metabolism (FDR = 3.26X10^-10^), protein deubiquitination (FDR = 0.0142), the ubiquitin-proteasome system (FDR = 0.0017), neurotransmitter transport (FDR = 0.011), collagen IV basement membrane organisation (FDR = 1.8X10^-4^), and trans-Golgi network vesicle budding (FDR = 2.4X10^-4^) (Figure 6D). Mitochondrial dysfunction is consistently reported in tauopathy ^49,50^, and multiple proteins important for regulating mitochondrial processes such as, VDAC1, CS, NDUFA10, NDUFS5, OGDHL, OXA1L, SUCLG1, ATAD3A, CHCHD6, CISD1, COX17, HSDL1, MICOS10, SLIRP, TIMM8B, and TOMM70A, were all reverted toward WT regulation following treatment with the ACR12 in K3 females (Figure 6D). Furthermore, several proteins important for autophagic (CTSB and SQSTM1/p62) and proteasomal (OTUB1, UCHL1, and USP46) protein degradation pathways were also reverted with treatment (Figure 6D). Additionally, GSK3β, a key regulator of tau phosphorylation, was decreased with ACR12 treatment (Figure 6D). Together, this suggests a potential role for ACR12 in promoting mitochondrial and proteasomal function and modulating pathological tau phosphorylation via downregulation of GSK3β.

To investigate a potential correlation between ACR12s effect on protein pathways and its direct interaction with tau, the 238 reverted proteins were overlayed with a tau interactome dataset compiling 12 tau interactome studies of human and rodent models of disease ^8^. Of the 238 proteins, 46 (19%) were identified as tau interactors. Furthermore, to ensure that the observed reversion effect is relevant to human disease, we validated our broader dataset of 1,412 proteins found to shift toward WT levels in ACR12-treated female K3 mice (see Figure 5I) against a human primary tauopathy, progressive supranuclear palsy (PSP), protein dataset identified via meta-proteomic analysis ^51^. Of our 1,412 proteins, 1,319 (93.4%) matched with orthologous proteins in the human PSP dataset. These proteins were identified to be enriched in a wide array of cellular and metabolic processes including, mitochondrial functions, cellular component organisation, and RNA splicing (Figure 6E). In summary, the functional enrichment and network analyses conducted here indicate potential roles for ACR12 in multiple protein pathways important for maintaining cellular and neuronal homeostasis, including but not limited to, RNA splicing, neuronal plasticity, synaptic processes, mitochondrial function, and protein degradation processes. Furthermore, ACR12-mediated reversion of these intracellular pathways aligns with the proteomic profile of the human PSP brain, suggesting that our findings have clinical relevance.

## DISCUSSION

Intracellular tau, rather than extracellular tau, makes up majority of tau in the brain^52^. And importantly, the toxic self-assembly of tau into aggregates occurs within neurons to trigger downstream neuronal and synaptic dysfunction^53–57^. Thus, targeting intracellular tau is a promising strategy for enhancing clinical outcomes. Previous studies have utilised tau-specific single-chain and single-domain intrabodies to target intracellular tau^58–61^, however, as tau in disease undergoes extensive posttranslational modifications, determining which specific epitope of tau to target is challenging^62^. We therefore aimed to target tau more broadly via neutralisation of the positively charged amino acids that make up the PRR and MTBD. To do this, we developed a novel peptibody, ACR12, comprised of a highly negatively charged peptide designed to electrostatically interact with the PRR and MTBD of tau, and fused to an inert scFv for downstream intraneuronal expression. *In vitro* binding assays confirmed that ACR12 engagement of tau was dependent on peptide charge, as peptibodies with less negative charge, ACR9 and ACR3, demonstrated reduced or no binding to tau. Furthermore, ACR12, via its scFv component, was expressed intracellularly *in vitro* and *in vivo*, and long-term treatment of K3 tau transgenic mice with ACR12 decreased pathogenic tau levels and restored brain proteostasis.

The marked reduction in total tau observed in ACR12-treated K3 mice suggests that ACR12, through interaction with tau, may facilitate its physiological degradation. This is supported by the proteomic analysis of ACR12-treated K3 brains which demonstrated a restoration of proteins with roles in ubiquitin-mediated proteolysis such as, UCHL1, Otub1, and sequestosome-1 (SQSTM1; also known as p62). For example, SQSTM1, in the context of tau, is required for the degradation of K63-polyubiquitinated tau lesions and, as such, its dysregulation worsens tau accumulation^63–65^. SQSTM1 also co-localises with tau NFTs in transgenic mice and in human Alzheimer’s disease (AD) tissue^7,64^. In addition to decreasing total tau levels in the K3 brain, ACR12 also reduced tau phosphorylation at pathogenic phosphorylation sites, T231, S235, and S202/T205. As these phospho-epitopes are within the PRR, this suggests that ACR12 may engage tau and directly prevent tau phosphorylation within this domain. This too is consistent with proteomic analyses revealing an ACR12-mediated decrease in levels of GSK3β, the predominant tau kinase which is elevated not only in K3 mice but also in other tauopathy mouse models^66,67^ and in human AD tissue^68,69^. GSK3β phosphorylates tau at multiple sites, including T231, S235, and S202/205^43,70^, and so, ACR12 may block GSK3β from interacting with tau, contributing to the observed reduction of cortical phosphorylated tau in K3 mice.

In addition to the effect of ACR12 treatment on tau pathology, ACR12-treated K3 mice also revealed a restoration of proteins involved in other neuronal pathways towards WT levels. Interestingly, ∼19% of ACR12-reverted proteins are known to directly interact with tau in disease. Other works employing affinity-purification mass spectrometry in tau-expressing cells, transgenic mice, human-derived iPSCs, and human brain tissue have identified various RNA-binding proteins^7,8,24,71^, regulators of ubiquitination and protein degradation^7,23,46,72^, and mitochondrial proteins^7,24,45,46^ as direct tau interactors. RNA processing and mitochondrial energy metabolism were the most prominent protein pathways to be restored by ACR12 treatment in K3 mice. RNA processing is known to be dysfunctional in tauopathy^8,22,48,57^ and importantly, RNA as a linear polyanion directly interacts with tau in disease to promote its aggregation^38,39,73^. Furthermore, several RNA-binding proteins (RBPs) such as HNRNPA1, HNRNPC, and HNRNPF, interact with tau in rodent models of disease and in human brain tissue^7,8,25,71,74,75^. This interaction is suggested to occur via electrostatic interaction of tau with RBP-bound RNA or positively charged regions within the RBPs themselves^76–78^. Thus, it is plausible that ACR12 may inhibit such interactions by neutralising the positively charged regions of tau. Similarly, mitochondrial dysfunction is well-established in tauopathies^50,79^, and overexpression of pathogenic tau has been shown to disrupt mitochondrial processes^80–82^. Tau interacts with a wide array of mitochondrial proteins^7,24,45,46^, several of which were restored toward WT level with ACR12 treatment including multiple subunits of the mitochondrial Complex I. Recent works revealed that tau can regulate mitochondrial reverse electron transport (RET), a process by which excess reactive oxygen species are produced^83^. Tau regulates RET via direct interaction with Complex I subunit, NDUFS3; this interaction requires phosphorylation of tau at the MTBD^83^. As such, ACR12 may inhibit this interaction via direct engagement with the MTBD and prevention of phosphorylation at this site. Together, the combination of analyses conducted in this study lead us to postulate that ACR12 may function by engaging the positively charged regions of tau to prevent pathological interactions of tau with other neuronal proteins.

There are, however, several limitations of the current work that must be considered. First, we were unable to investigate the effect of ACR12 on tau aggregation in this study as the K3 mice at study endpoint did not have sufficient insoluble tau to quantify. Future work in an alternate tau transgenic mouse model with greater pathology will allow us to investigate whether ACR12 can inhibit downstream tau aggregation. Second, as ACR12 binds to tau via electrostatic interaction, it is possible that ACR12 may interact with other positively charge proteins in the neuron. We therefore cannot conclude that all therapeutic effects observed here are mediated by the tau/ACR12 interaction. Finally, a significant sex effect was observed, wherein female and male K3 mice displayed distinct proteomic alterations relative to WT mice. Furthermore, the effect of ACR12 treatment was more pronounced in female mice than males and thus, downstream proteomic analyses were restricted to female mice. Importantly, sex differences are not unique to this study and have been observed previously in the K3 mouse model and in other tau transgenic mice^84–86^, warranting future studies that investigate the mechanisms driving this sex effect.

In conclusion, this study suggests a key role for the exposed positively charged residues of tau in pathogenesis. As such, therapeutics designed to neutralise these regions of tau, such as ACR12, may be effective in ameliorating tau pathology. Moreover, this work introduces the peptide-scFv peptibody as a novel therapeutic modality for targeting intracellular tau that could be applied to other therapeutic peptides to enhance intracellular activity.

## METHODS

### Antibodies

Primary antibodies used for western blotting (WB), immunofluorescence (IF), and immunoprecipitation (IP): α-Flag (WB: 1:1,000, IF: 1:800; Thermo Fisher Scientific), β-actin (WB: 1:1,000; Thermo Fisher Scientific), α-GFP (IP: 1:200; Abcam), α-tau A0024 (polyclonal tau antibody) (WB: 1:1,000; Dako), MAP2 (IF: 1:500, Abcam), Tau-5 (specific for tau aa210-241, WB: 1:1,000; Thermo Fisher Scientific), AT8 (phospho-tau S202/T205, IF: 1:500; Thermo Fisher Scientific), AT180 (phospho-tau T231, IF: 1:500; Thermo Fisher Scientific), RN235 (phospho-tau S235, IF: 1:500; Sigma) The secondary antibodies used were as follows: IRDye 680RD goat anti-mouse IgG (WB: 1:10,000; LI-COR), IRDye 800CW goat anti-rabbit IgG (WB: 1:10,000; LI-COR), goat anti-mouse Alexa Fluor 594 (IF: 1:500; Thermo Fisher Scientific), goat anti-mouse Alexa Fluor 488 (IF: 1:500; Thermo Fisher Scientific), donkey anti-chicken Alexa Fluor 555 (IF: 1:500; Thermo Fisher Scientific).

### Production of peptibody constructs

Synthetic genes encoding ACR3, ACR9, and ACR12 peptibodies were generated, each comprising of a unique, positively charged peptide fused to the C-terminal end of a neutral scFv isolated as a control antibody from a Tomlinson I+J library as described previously^84^. Synthetic genes were cloned into the AAV mammalian expression plasmid, pAM-CBA-pI-WPRE-BGHpA (provided by Matthias Klugmann UNSW) and sequence confirmed via the Australian Genome Research Facility (AGRF). Recombinant ACR peptibodies and tau proteins were produced as described previously^45^.

### ELISAs

The binding specificity and EC_50_ of ACR peptidbodies were determined using an enzyme-linked immunosorbent assay (ELISA). Briefly, Immuno 96 MicroWell plates (Nunc) were coated with 10μg/mL recombinant 2N4R full-length human tau (hTau441) or K18 tau peptide (MTBD region), followed by blocking with 3% BSA in PBS. Wells were then incubated with 10μM ACR3, ACR9, or ACR12 recombinant protein. Bound ACRs were detected using α-Flag/His antibodies followed by anti-rabbit IgG-HRP conjugate (1:5000), and incubation with TMB substrate (Sigma-Aldrich); absorbance was measured at 450nm. Measurements were performed in triplicate and analysed following subtraction of PBS only absorbance. For EC_50,_ plates serial dilutions of ACR proteins were applied.

### ThT assay

Recombinant hTau441 protein (5μM) was incubated with 50μM ThT, 1mM DTT, and 10μM ACR9 or ACR12, or heparin (1.25 μM) as a positive control for tau fibrillization, over 72h at 37*C with shaking prior to each absorbance reading. ThT fluorescence measured every 20min with excitation and emission wavelengths of 440 and 480nm, respectively. Averaged curves were generated from triplicate measurements.

### Cell culture

SH-SY5Y human neuroblastoma cells that overexpress full-length human tau tagged with GFP at the C-terminus (Tau-GFP cells) were cultured in Dulbecco’s modified Eagle’s media (Gibco), supplemented with 10% (*v/v*) foetal calf serum (Gibco), 1% (*v/v*) penicillin-streptomycin (Gibco), and maintained with 6mg/mL blasticidin (Thermo Fisher Scientific) at 37°C with 5% CO_2_ and ≥ 80% relative humidity.

### Cell transfections and lysis

Tau-GFP cells were transfected with ACR peptibody plasmid DNA using Lipofectamine 2000 (Thermo Fisher Scientific) as per manufacturer’s protocol. Cells were collected and lysed 24h post-transfection in 1X RIPA buffer supplemented with cOmplete Protease Inhibitor cocktail tablet with EDTA (Roche) for 30min on ice. Samples were centrifuged at 10,000*xg* for 10min at 4°C, supernatants collected, and total protein assessed using a BCA assay (Thermo Fisher Scientific).

### Immunoblotting of transfected cell lysates

Cell lysates collected from ACR-transfected Tau-GFP cells were diluted with 4X NuPAGE LDS Sample Buffer (Thermo Fisher Scientific) and 10X NuPAGE Sample Reducing Agent (Thermo Fisher Scientific), electrophoresed on 4-12% Bis-Tris NuPAGE gels (Thermo Fisher Scientific), and transferred to a Low Fluorescence PVDF membrane using the iBlot 2 Dry Blotting System (Thermo Fisher Scientific). Membranes were stained with REVERT 700 Total Protein Stain according to manufacturer’s instructions (LI-COR) and imaged at 700nm with the Odyssey Scanner (LI-COR). Membranes were de-stained and blocked for 1h in 1X Blocker FL Fluorescent Blocking Buffer (Thermo Fisher Scientific) and probed with primary antibody solution overnight at 4°C. Membranes were imaged and fluorescence quantified using Image Studio Lite (LI-COR).

### Immunofluorescence and microscopy of transfected cells

Tau-GFP cells plated onto glass coverslips were transfected with ACR peptibody plasmid DNA as previously described prior to fixation, permeabilization, and blocking with 5% goat serum 1% bovine serum albumin in PBS for 1h. Cells were then incubated with α-Flag antibody overnight at 4°C, washed, probed with goat anti-mouse Alexa Fluor594 (Thermo Fisher Scientific), and nuclei labelled with DAPI nuclear stain (4’, 6-diamidino-2-phenylindole; Sigma-Aldrich) for 1.5h in the dark. Coverslips were then mounted onto slides with Fluorescent Mounting Media (Agilent) and imaged using the Crest Spinning Disk (Nikon) under the 60x oil objective; the system was controlled by NIS Elements acquisition software. Confocal *z*-stacks were imported into the ImageJ software and exposure adjusted. Co-localisation analyses were conducted on the entire *z*-stack for single cells using the Coloc2 plugin, calculating Pearson’s correlation coefficient. Pearson’s coefficient (*r*) measures the degree to which signal intensities in two channels are linearly correlated to each other, by which *r* = 1 is perfect positive correlation and *r* = 0 is no correlation. Transfection efficiency was determined in IMARIS analysis software using the Spots creation tool to quantify transfected (Flag-positive) and total cells. Estimated XY diameter for Spot detection was 5μm.

### Co-immunoprecipitation assay

Tau-GFP cells co-transfected with ACR9 or ACR12 were lysed in 1X Triton-X Lysis Buffer (20mM Tris-HCl pH 7.4, 10mM potassium chloride, 10mM magnesium chloride, 2mM EDTA, 10% glycerol (*v/v*), 1% Triton X-100 (*v/v*)). An aliquot of each lysate was collected as ‘input’ and stored at −20°C until future use. Remaining lysates were made up to 500μL with lysis buffer and incubated with α-GFP antibody (Abcam) overnight at 4°C with constant rotation. Protein G Agarose beads (Roche) were then applied for 1h with rotation, and sample centrifuged for 2min at 5000*xg*. Supernatants were discarded and beads washed thrice with lysis buffer prior to resuspension in 4X NuPAGE LDS Sample Buffer. To elute, beads were boiled for 10min at 95°C and centrifuged for 3min at 5000*xg;* supernatants were collected (‘IP’). Input and IP samples were then electrophoresed and immunoblotted as previously described.

### Animals

Animal experiments were conducted following the guidelines of the Australian Code of Practice for the Care and Use of Animals for Scientific Purposes and approved by the Florey Animal Ethics Committee (AEC#23-001-FINMH). For AAV ACR9 and ACR12 expression and treatment studies, the K3 mouse model expressing human 1N4R tau with a K369I mutation via mThy1.2 promoter was used ^41^ (a generous gift from Professor Jürgen Götz, University of Queensland). Experimental K3 mice were generated by breeding male K3 mice with wild-type (WT) females. Age-matched WT littermates were used as healthy controls in all behavioural experiments. All mice were housed in pathogen-free cages, maintained on a 12h light/dark cycle and had constant access to food and water; mice were weighed weekly from four-weeks-old until study endpoint.

### AAV-PHP.eB ACR vector preparation and tail vein injections

The AAV-PHP.eB ACR9 and ACR12 vectors were produced by the UQ Viral Vector Core (University of Queensland), using peptibody plasmids prepared as described.

For AAV ACR9 and ACR12 expression studies, five-week-old female and male WT and K3 mice were injected with saline *(n=*2 WT, *n=*2 K3) or AAV-PHP.eB ACR9 (*n=*2 K3) or AAV-PHP.eB ACR12 (*n=*2 WT, *n=*2 K3) intravenously into the lateral tail vein. A dose of 2.5 × 10^10^vg/mouse was used. For treatment and behavioural studies, K3 mice were randomly assigned to one of the following treatment groups: saline (Sham; *n=*6♀, *n=*6♂), AAV-PHP.eB ACR9 (*n*=6♀, *n*=6♂), or AAV-PHP.eB ACR12 (*n*=6♀, *n*=7♂). At five-weeks-old mice were injected intravenously as described above.

### Behavioural tests

Mice were tested fortnightly on the Rotarod to track motor function, once prior to injection and then up until study endpoint. Mice were placed on the Rotarod and assessed using linear acceleration mode (4 to 40rpm) over 90s, with a total test time of 180s (90s at 40rpm). Latency to fall was recorded as the mean time a mouse spends on the Rotarod out of five attempts. At 20-weeks-old, mice underwent Digigait analyses. The front and hind paws of each mouse were coloured with red food colouring and each animal was placed onto the Digigait treadmill set to 15cm/s. Run was completed following recording of 10-12 steps using the Digigait software (Mouse Specifics Inc.). For both behavioural tests, light levels in the testing room were kept constant at 60% (∼70lx) and mice were habituated to the testing environment for 30min prior to testing.

### Tissue collection and processing

Mice were anaesthetised with an overdose of sodium pentobarbitone and transcardially perfused with PBS. Brains were collected and separated into hemispheres. Right hemispheres, as well as peripheral organs, were snap-frozen in liquid nitrogen and stored at – 80°C until processing. Left hemispheres were immersion-fixed in 4% PFA for 24h at 4°C, washed twice with PBS, and submerged in 30% sucrose 0.1% sodium azide solution for at least 48h at 4°C prior to cryo-sectioning. Sagittal cryo-sections at 20μm thickness were obtained in a cryostat and stored at –80°C until staining.

Snap-frozen hemispheres were weighed and homogenised using a gentle method optimized to maintain protein-protein interactions ^87^. Briefly, hemispheres were pulverised with 10 hammer blows prior to dounce homogenisation on ice (∼25 strokes) in 5ml/g low salt homogenisation buffer (50mM HEPEs pH7.0, 250mM sucrose, 1mM EDTA) supplemented with protease and phosphatase inhibitors (Millipore Sigma). Total protein concentration of homogenates was determined via BCA assay. 15μg of collected homogenates were electrophoresed, immunoblotted, and analysed as previously described to assess ACR peptibody expression and tau protein levels. A fraction of homogenates was aliquoted for use in proteomics as described below.

### Immunofluorescence and immunohistochemistry of mouse brain sections

Slides containing collected mouse brain cryo-sections were placed into a humidity chamber and washed thrice with PBS and blocked in 10% goat serum, 1% BSA, 0.5% triton-X TBS for 2h at room temperature. Slides were then washed twice with PBS prior to incubation in primary antibody solution overnight at 4°C. The next day, slides were washed and sections stained with DAPI nuclear stain and secondary antibody solution for 1.5h at room temperature in the dark. Slides were washed a final time and coverslips mounted with Fluorescent Mounting Medium. All images (20x) were taken using the Axioscan7 (Zeiss) microscope under control of the ZEN blue acquisition software.

All immunofluorescence staining quantification was performed blinded and using ImageJ. Briefly, secondary only control sections were used to set the background threshold for each slide. Two or three sections per mouse were thresholded according to control, and percentage (%) area AT180 (phospho-tau T231), AT8 (phospho-tau S202/T205), and RN235 (phospho-tau S235) tau positive staining in the cortex was quantified using the ‘Analyse Particles’ function, wherein particle size was set to 50μm and circularity to 0.05-1.00 to isolate appropriate structures ^88^. For all analyses, tissue that was damaged or had high background staining due to autofluorescence was excluded.

### LC-MS/MS

Approximately 100µL of each snap-frozen brain homogenate was added to SDS (2% w/v) and TCEP at 10mM, prior to heating for 10min at 85°C. Samples were then cooled to <25°C and incubated at 23°C for 30min with 20mM iodoacetamide. Samples were precipitated using the chloroform-methanol method, air-dried, and redissolved in 20µL 7.8M Urea with 100mM HEPES pH 8.0 and 5μg LysC endoproteinse (FUJIFILM Wako). LysC digestion was for 6h at 28°C. Samples were diluted 8-fold with 100mM HEPES pH 8.0 and two trypsin digestions were done for 8h at 28°C, each with 5μg trypsin (TrypZean, Sigma). The approximate amount of protein was determined by UV absorption at 280nm using a Nanophotometer N60 (Implen). Approximately 300μg tryptic peptides from each sample were labelled with TMTpro (18plex) (Thermo Fisher Scientific).

After confirming efficient labelling, each sample set was mixed into 16plexs and de-salted using solid phase extraction cartridges (Sep-Pak, Vac 3 cc 200 mg tC18, Waters). Hydrophilic ion liquid chromatography (HILIC) was performed on a Vanquish Neo HPLC system (Thermo Fisher Scientific) with a 250mm long and 1mm inside diameter TSKgel Amide-80 column (Tosoh Biosciences). The HILIC gradient used a solution of 99.9% acetonitrile, 0.1% TFA (buffer A), and a solution of 0.1% TFA (buffer B). The sample was loaded onto the system with a flow rate was 50μl/min in 90% buffer B. Fractions were collected into a 96-well plate using an FC204 fraction collector (Gilson) every 60s and monitored by absorbance of UV light at 214nm. The UV signal was used to combine selected fractions into similar amounts of peptide. Fractions were dried and reconstituted in 0.1% formic acid for LC-MS/MS analysis.

The LC-MS/MS was performed using a Vanquish Neo UHPLC system and Astral Orbitrap mass spectrometer (Thermo Fisher Scientific). Each HILIC fraction was loaded directly onto an in-house 300mm long 0.075mm inside diameter column packed with ReproSil Pur C18 AQ 1.9μm resin (Dr Maisch, Germany). The column was heated to 50°C using a column oven (PRSO-V2, Sonation Lab Solutions) integrated with the nano flex ion source with an electrospray operating a 2.3kV. The S lens radio frequency level was 50 and capillary temperature 280°C.

The liquid chromatography used buffer A (0.1% formic acid) and buffer B (0.1% formic acid, 80% acetonitrile). After loading sample in buffer A, the gradient, at 300nL/min, was from 7% buffer B to 27% buffer B in 92min, to 35% buffer B in 2min, to 99% buffer B in 1min, and held at 99% buffer B for 3min; MS acquisition was 100min. All samples and fractions were analysed using data-dependent acquisition. The MS scans in the orbitrap were at a resolution of 120,000 with automatic gain control target set to standard for a maximum ion time of 50ms from 375 to 1500m/z. The MS/MS scans in the Astral analyser used standard automatic gain control target with a maximum ion time of 10ms. Up to 50 MS/MS scans per cycle were allowed with an intensity threshold of 50,000counts. The isolation window was 0.7m/z, the range was 120-1400 and the normalized collision energy was 35. Charge state from +2 to +6 were allowed and dynamic exclusion for 18s was performed after 2 occurrences within 30s.

Raw LC-MS/MS data was processed with MaxQuant v2.6.3.0. Variable modifications were oxidation (M), acetyl (protein N-terminus), deamidation (N and Q) and phosphorylation (S, T, and Y). Carbamidomethyl (C) was a fixed modification. Digestion was set to trypsin/P with a maximum of 3 missed cleavages. The TMTpro correction factors were entered. The *Mus musculus* reference proteome was used with canonical and isoform sequences (downloaded on Dec 23, 2024 with 54,742 entries and 21,757 genes). A fasta file containing the human tau isoform E was also used, as well as the inbuilt MaxQuant contaminants fasta file. The minimum peptide length was six; second peptides and dependent peptides searches were enabled; peptide spectrum matching and protein false discovery rates were set at 1%.

### Processing of MaxQuant proteome data

Additional processing of the MaxQuant spectra output “evidence.txt” and “proteinGroups.txt” files for the proteome was performed in R v4.3.2. Protein IDs were annotated to gene symbols as previously described ^89–91^, using the same fasta file employed for the MaxQuant spectra search; only log_2_ transformed intensity values from complete data (no missing values for any replicates) were retained for further analysis. Data was quantile normalised ^92^ and then underwent surrogate variable analysis for removal of unwanted sources of variation using a predefined model describing the conditions. Mixed control samples were removed prior to surrogate variable analysis ^93^. The selection of k, number of surrogate variable required, to remove batch effects and recover sample grouping was guided by hierarchical clustering and PCA of 1) the combined male and female samples across batches and 2) resultant clustering of each batch of male and female samples. For statistical analysis, limma v3.58.1, was utilised for fitting generalized linear models with Bayes shrinkage for differential abundance of proteins ^94^. P-values obtained from the moderated t test were then corrected for multiple hypotheses with the Benjamini and Hochberg (FDR) method ^95^. DEPs were defined as those with p < 0.05 after FDR correction.

### Cell specificity analysis

Cell specificity analyses was performed using single-cell RNA-seq (scRNAseq) gene expression data and cell cluster annotations from The Human Protein Atlas (data downloaded on 19/11/2025 from: https://www.proteinatlas.org/download/tsv/rna_single_cell_cluster.tsv.zip)^47^. Data was imported into Rstudio Server (R version 4.3.2) and filtered by “Tissue” type as “brain”. 19 primary cell type annotations (“Cell type”) were used for analysis by summing the normalised counts per million (nCPM) for cells annotated to multiple sub-cluster annotations and then log2 transforming the nCPM (with a psuedocount addition of 1 to account for zero counts). Human gene symbols were remapped to homologous mouse genes using “Orthology.eg.db” version 3.18.0. Genes encoding the intersecting DEPs found in female only K3vWT and K3 ACR12vK3 Sham comparisons was used at the input gene set for cell specificity analysis and the log2 nCPM expression for each gene determined from the scRNAseq data. A boxplot of the log2 nCPM expression for each cell type was generated using ggplot2 v4.0.0 (ordered by median expression) and a heatmap of z-score transformed (row-wise) logCPM expression of genes generated using ComplexHeatmap v2.18.0 (clustering rows and columns, with columns k-means clustered at k-3).

### Protein functional enrichment and network analyses

Protein enrichment analyses were performed using STRING (Version 12.0), employing the following ontology databases: *Mus Musculus,* GO Biological Processes, and GO Cellular Components ^96^. Intersecting DEPs found in female only K3vWT and K3 ACR12vK3 Sham comparisons, representative of proteins that were both altered in disease and responsive to treatment, were separated by change direction (increasing or decreasing in abundance), and inputted into STRING. Enrichment results were imported into RStudio v4.4.2 and bubble plots generated using ggplot2 v3.5.1 and patchwork v1.3.0 packages to visualise the top 10 (by FDR) enriched terms. For PPI analysis, network data from STRING was imported into Cytoscape v3.10.3 and further analysed using the MCODE algorithm to identify densely connected protein clusters ^97^.

### Statistical analysis

All statistical analyses (excluding proteomic data) were performed using GraphPad Prism 10 software. Statistical significance between experimental groups was analysed using either one-way ANOVA followed by Tukey’s multiple comparisons test or unpaired *t*-test, where *P* < 0.05 was considered significant. Outliers were removed where required using the ROUT (Q=1) method. All values are reported as mean ± standard error of the mean.

## DATA AVAILABILITY

The mass spectrometry proteomics data have been deposited to the ProteomeXchange Consortium via the PRIDE partner repository with the dataset identifier PXD066615 and 10.6019/PXD066615. Other data supporting the findings of this study are available from the corresponding author, upon reasonable request.

## ACKNOWLEDGEMENTS

The authors thank Florey Core Animal Services for their support in this work. The authors gratefully acknowledge the Biological Optical Microscopy Platform and the Florey Neuroscience Microscopy Facility for their support and assistance in this work. We thank Celeste Mawal, Adam Southon, and Scarlett Parker for managing the lab. We thank Alicia Yong, for assistance in tissue processing, as well as Madeleine Lester and Anne Miller for assistance in recombinant protein production. We acknowledge the traditional custodians of the land on which this work was conducted, the Wurundjeri people of the Kulin nation.

## AUTHOR CONTRIBUTIONS

Conceptualization: ACC, ED, RM; Methodology: ACC, AJW, MG, RMN; Investigation: ACC, TK, AJW, MG, RMN; Formal analysis: ACC, TK, AJW, ED, RMN; Visualization: ACC, TK, ED, RMN; Writing – original draft: ACC, RMN; Writing – review & editing: ACC, ED, RMN; Funding acquisition: RMN; Supervision: ED, RMN

